# Collateral fitness effects of mutation are not commonly caused by protein misfolding

**DOI:** 10.1101/2025.09.12.675869

**Authors:** N. Quan, Y. Eguchi, A. Brown, K Geiler-Samerotte

## Abstract

Mutations in coding sequences are often assumed to harm cells by destabilizing proteins and creating toxic misfolded species. Here we directly test how fitness scales with predicted folding stability. Using deep mutational scanning of a gratuitously expressed protein in *S. cerevisiae* (≈2,000 YFP single-amino-acid variants) and meta-analyses of seven additional scans of gratuitous proteins in yeast and *E. coli*, we find that collateral fitness effects, costs that arise independently of protein function, do not correlate with predicted destabilization (ΔΔG). Even variants predicted or biochemically shown to misfold frequently had no measurable collateral cost. In contrast, across matched datasets where the same proteins were required for growth, predicted destabilization strongly tracked primary fitness costs, and this association intensified as functional demand increased. These conclusions were robust to multiple stability predictors and to competitive fitness assays with high sensitivity. Together, our results indicate that misfolding is not a common driver of collateral fitness costs, whereas it often underlies primary costs when function matters. These findings overturn the long-standing assumption that misfolding universally drives the collateral costs of mutation, reframing misfolded proteins as only one piece of a broader puzzle and opening the way to identify alternative cellular vulnerabilities that shape evolution, disease, and aging.

## Introduction

Misfolded proteins are often described as being harmful to cells (Geiler-Samerotte et al. 2011; Karapanagioti et al. 2024; Brezic et al. 2025; Wu et al. 2025) and are associated with diverse neurological diseases (Hartl 2017). Previous work has shown that misfolded proteins can puncture cell membranes (Gonzalez-Garcia et al. 2021), damage the mitochondria (Abramov et al. 2020; Alqahtani et al. 2023), provoke stress response systems (Geiler-Samerotte et al. 2011, 2013), siphon proteostasis resources (Gidalevitz et al. 2009), and mis-interact with other cellular components (Yang et al. 2012). Since the majority of coding mutations destabilize protein folding (Pakula and Sauer 1989), mutations may represent a steady source of problematic misfolded molecules. Consistent with this view, mutations appear to be purged by natural selection in accordance with the number of misfolded proteins they stand to produce, accumulating at slower rates in highly expressed genes (Drummond et al. 2005; Drummond and Wilke 2008). The resulting correlation between a protein’s evolutionary rate and its expression level is one of the most pervasive patterns across the tree of life (Pál et al. 2001; Zhang and Yang 2015). In addition to informing theories about how proteins evolve, the idea that misfolded proteins tend to harm cells has also inspired cancer treatment models that exploit weaknesses created by a tumor’s elevated mutation rates (McFarland et al. 2013, 2017; Tilk et al. 2024), as well as anti-aging strategies that upregulate proteostasis machinery to alleviate the harm caused by widespread misfolding of cellular proteins (Morimoto and Cuervo 2014). Despite a few examples of misfolded proteins that do not appear to be toxic (Plata et al. 2010; Kafri et al. 2016), the dominant hypothesis, one which has influenced disparate fields including neurobiology (Soto and Pritzkow 2018), evolution (Drummond and Wilke 2008), cancer (Tilk et al. 2024), aging (Drummond and Wilke 2008; Santra et al. 2019), cell biology (Oromendia et al. 2012) and climate biology (Bongioanni et al. 2021; Sisodiya et al. 2024), is that misfolded proteins generally tend to be harmful.

Here, we sought to measure how the harmfulness of misfolded proteins scales with their abundance, an open question put forth in previous work (Echave and Wilke 2017; Eguchi et al. 2019). In the model eukaryote, *S. cerevisiae*, we generated a large dataset using deep mutational scanning of a ‘gratuitous’ protein (i.e. one that confers no functional benefit to an organism). Specifically, we created over 2,000 single mutant variants of yellow fluorescent protein (YFP) and competed them against a strain expressing the unmutated version at the same level. Previous studies have shown that mutations to gratuitous proteins can impact fitness. Such effects, termed “collateral fitness effects,” are distinguished from fitness consequences due to loss of function, known as “primary fitness effects” (Mehlhoff et al. 2020) (**Fig 1A**). In line with this framework, we found that many mutations to our gratuitous protein reduced fitness, supporting prior observations that collateral fitness effects are common (Geiler-Samerotte et al. 2011; Mehlhoff et al. 2020). While collateral costs can arise for many reasons (Kintaka et al. 2016; Eguchi et al. 2018; Mehlhoff and Ostermeier 2023), they are most often attributed to a gain of toxic misfolded proteins (Geiler-Samerotte et al. 2011, 2013; Serohijos et al. 2012; Oromendia et al. 2012; Mehlhoff et al. 2020; Wu et al. 2022). However, the collateral fitness costs of mutations to YFP does not correlate with their predicted effects on protein stability. Strikingly, even mutations predicted to cause substantial destabilization often had no detectable fitness cost, while some of the most costly mutations had no predicted impact on folding. To evaluate the generality of this pattern, we compiled data from an additional seven deep mutational scans of gratuitously expressed proteins (Stiffler et al. 2015; Wu et al. 2022; Mehlhoff and Ostermeier 2023). Across all datasets, regardless of the protein, organism, or experimental design, we observed the same phenomenon: the predicted extent of misfolding showed no relationship to fitness.

**Figure 1:**
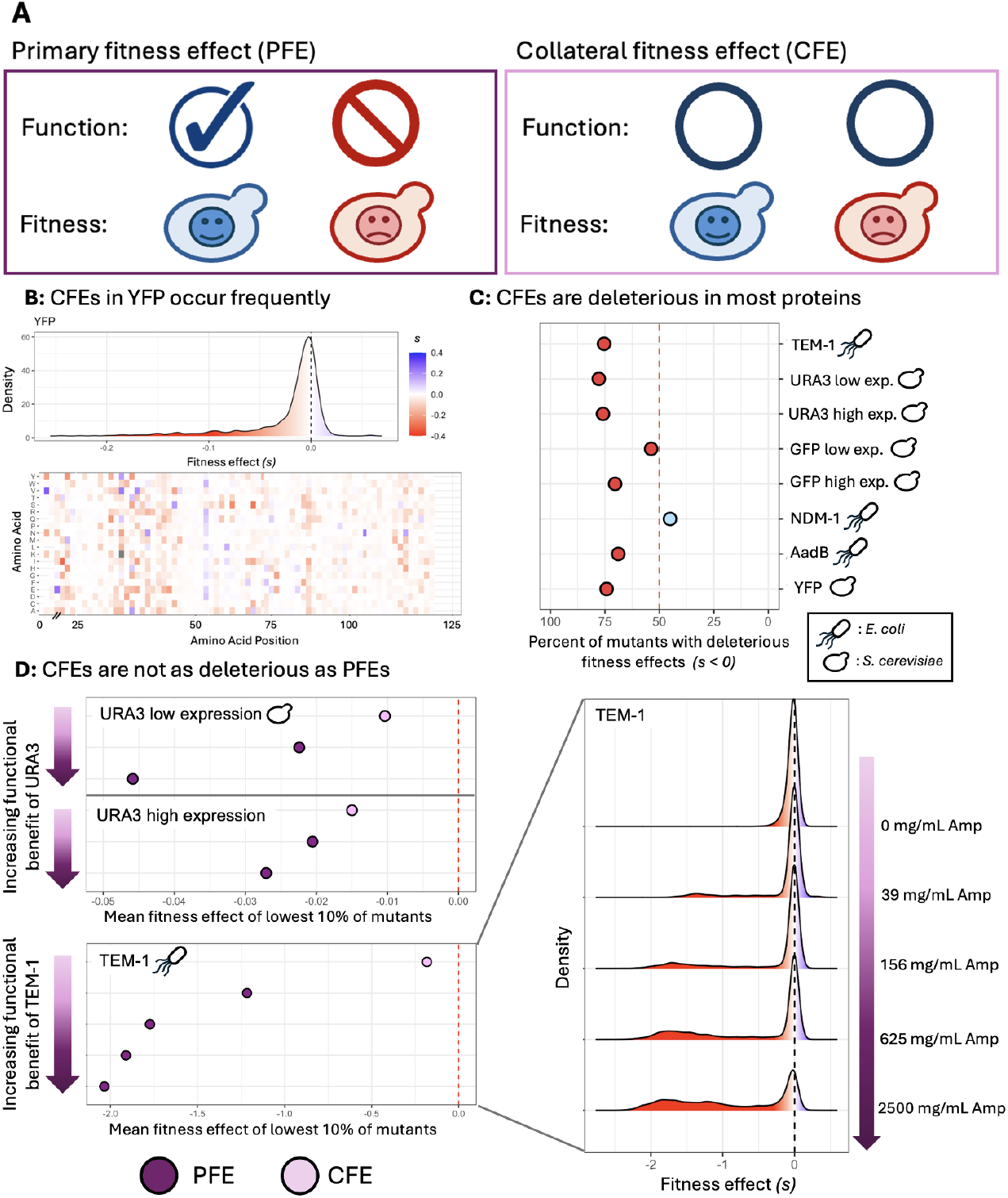
Collateral fitness costs are prevalent and less severe than primary fitness costs. **(A)** Primary fitness effects result from loss of protein function. Collateral fitness effects arise independently of protein function. **(B)** A deep mutational scan of 1,995 amino acid positions in yellow fluorescent protein (YFP) reveals pervasive collateral fitness costs in yeast cells. Fitness effects are red when deleterious, represent the average across two replicates, and are measured relative to a strain expressing wildtype YFP. Heat map shows the location and fitness effect of each amino acid substitution. **(C)** Collateral fitness effects tend to be deleterious across 8 different deep mutational scans in yeast and E. coli. **(D)** Primary fitness costs tend to be larger than collateral ones. The horizontal axis displays the average fitness of the least fit 10% of mutants, which is lower for PFEs (dark purple) relative to CFEs (light purple). The functional benefit of URA3 (involved in uracil production) was increased by removing uracil from media (Wu et al. 2022). The functional benefit of TEM1 (ampicillin degradation) was increased by adding ampicillin to the media (Stiffler et al. 2015). In both cases, doing so increases the deleterious fitness consequences of mutations to the protein of interest.

One possible explanation is that our predictions of folding disruption, estimated following previous work, using MutateX (Tokuriki et al. 2007; Pancotti et al. 2022; Tiberti et al. 2022), are flawed. To address this, we turned to eight matched datasets from the same mutant libraries, but in conditions where the proteins were required for growth and thus no longer gratuitous. In these non-gratuitous settings, predicted misfolding and fitness loss were tightly correlated. Mutations predicted to destabilize the folding of beneficial proteins consistently reduced fitness, and there were no cases where highly misfolded variants escaped fitness costs. These findings, and additional data from other folding prediction algorithms, suggest that the observed independence of collateral fitness costs and protein misfolding is not a technical artifact.

Another possible reason for the observed lack of connection between misfolding and collateral fitness costs could be that the latter are too small to detect in laboratory experiments. However, previous work was able to quantify such costs by overexpressing a handful of misfolded YFP variants using a smoothly inducible galactose promoter (Geiler-Samerotte et al. 2011). Here, we expressed YFP to even higher levels by using a smoothly inducible ATC promoter (Azizoglu et al. 2021; Quan et al. 2023). Despite this, we still could not detect significant fitness costs for most destabilized YFP variants. Further, all of the fitness measurements we obtained from the literature were measured using competitive fitness assays, which are among the most sensitive approaches available for detecting small fitness effects (Mehlhoff et al. 2020; Wu et al. 2022; Mehlhoff and Ostermeier 2023; Kinsler et al. 2023). Despite these highly powered experimental designs, we still failed to detect consistent collateral fitness costs associated with protein misfolding. However, we do observe many collateral fitness costs, some as large as 20%, in variants that are not predicted to misfold. These observations are consistent with previous studies suggesting that the toxicity of protein misfolding is not always the strongest contributor to the collateral fitness costs of mutation (Kintaka et al. 2016; Eguchi et al. 2018; Mehlhoff and Ostermeier 2023).

Together, our results support two related conclusions. First, our work suggests that protein misfolding does not always lead to toxicity. By pinpointing misfolded proteins that are unexpectedly well-tolerated, our dataset provides a foundation for uncovering the molecular features that distinguish harmful misfolded proteins from those that are benign. Second, our work suggests that many collateral fitness effects arise from mechanisms other than misfolding. This finding motivates a deeper investigation into the major sources of collateral fitness costs that arise independently of folding defects. Understanding the molecular basis of these costs is critical for predicting evolutionary trajectories, illuminating basic aspects of cell biology, and developing strategies to treat mutation-driven diseases like cancer.

## Results & Discussion

### Many mutations to gratuitously expressed proteins decrease fitness, suggesting that collateral fitness defects are common

The first step to studying collateral fitness costs is to distinguish these costs from those associated with loss of functional proteins. This is often done by studying mutations to proteins that have no beneficial functions in the conditions where they are expressed. Previous work has shown that mutations to such gratuitously expressed proteins can indeed impose costs relative to wildtype versions of the protein expressed at the same level (Geiler-Samerotte et al. 2011; Mehlhoff et al. 2020; Wu et al. 2022). These costs have been called “collateral fitness costs” (Mehlhoff et al. 2020) to distinguish them from primary fitness costs that are due to changes in a protein’s function (**Fig 1A**). The prevalence and magnitude of collateral fitness costs range across proteins (Mehlhoff and Ostermeier 2023), conditions and expression levels (Wu et al. 2022), with up to 42% of nonsynonymous mutations to a given protein resulting in collateral costs (Mehlhoff et al. 2020). While collateral costs can arise for many reasons (Mehlhoff and Ostermeier 2023), they are most often attributed to a gain of toxic misfolded proteins (Geiler-Samerotte et al. 2011, 2013; Serohijos et al. 2012; Oromendia et al. 2012; Mehlhoff et al. 2020; Wu et al. 2022).

Here, we conducted a deep mutational scan for yellow fluorescent protein (**Fig 1B; Supplementary Table 3, Supplementary Table 6**), and compiled additional scans from previous work, resulting in 8 datasets describing the collateral fitness costs of amino acid substitutions spanning different proteins, organisms and expression systems (**Fig 1C**) (Stiffler et al. 2015; Mehlhoff et al. 2020; Wu et al. 2022; Mehlhoff and Ostermeier 2023). When primary fitness costs are absent, most substitutions have negative fitness effects (s < 0) (**Fig 1C**), consistent with previous findings suggesting that collateral fitness effects are common and are deleterious (Geiler-Samerotte et al. 2011; Mehlhoff et al. 2020; Wu et al. 2022; Mehlhoff and Ostermeier 2023).

We also compiled matched mutational scans for 3 of the above proteins in multiple conditions where expression is no longer gratuitous (Stiffler et al. 2015; Wu et al. 2022) (**Fig 1D**). When primary fitness costs are present, amino acid substitutions tend to have larger costs (**Fig 1D**; dark purple points move farther to the left). These costs increase as the function of the protein becomes more important for survival. For example, TEM1 is an enzyme that renders ampicillin ineffective. As ampicillin concentration increases, the fitness costs of mutations to TEM1 tend to increase (**Fig 1D**; the red peak becomes taller and broader with increasing ampicillin concentration). Next we asked whether any of these costs, collateral or primary, could be predicted by the extent of protein misfolding.

### Effects on protein folding predict primary but not collateral fitness costs

Previous work suggests that the toxicity of misfolded proteins provokes a collateral fitness cost that increases with the extent of misfolding (Drummond and Wilke 2008; Geiler-Samerotte et al. 2011). Therefore, we expected fitness to decrease with each mutation’s effect on protein stability (**Fig 2A**). This is not what we observed for collateral fitness costs in most of the 8 proteins we study (**Fig 2B; Fig S1**). A mutation’s effect on stability, as inferred from stability prediction algorithms such as MutateX (Tiberti et al. 2022), cannot predict collateral fitness costs (**Fig 2B**; **Fig S1**).

**Figure 2:**
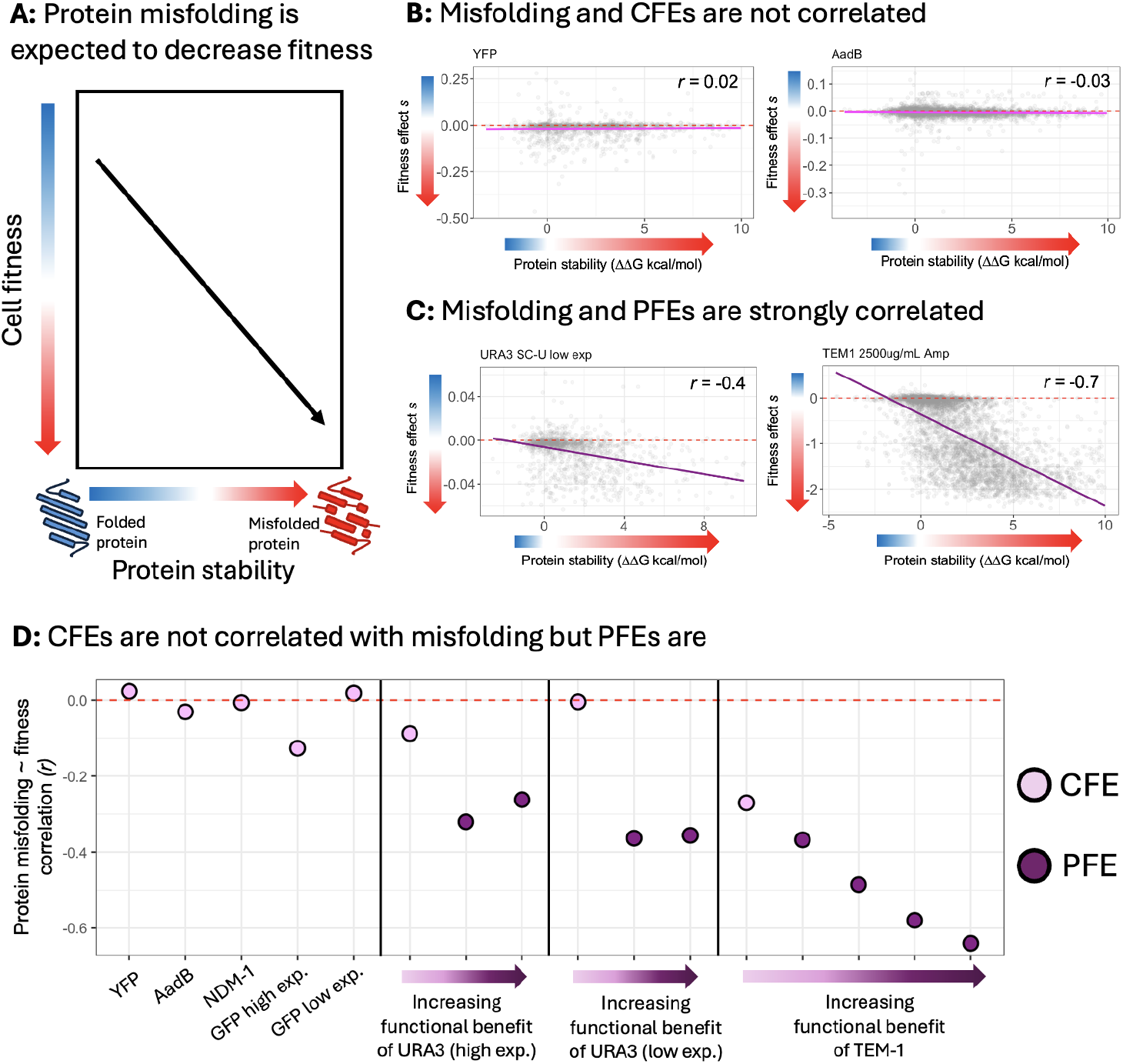
Collateral fitness costs are not correlated with protein misfolding. **(A)** Toy model showing expectation that mutations associated with protein misfolding should tend to decrease fitness. **(B)** Two of eight graphs showing that mutations associated with protein misfolding do not tend to result in collateral fitness effects (see others in **figure S1**). Each point represents a single amino-acid substitution. The vertical axis depicts its fitness effect. The horizontal axis depicts its predicted effect on protein stability using MutateX. Pink lines display near-zero Pearson correlation between fitness and stability. **(C)** Two of eight graphs showing that mutations associated with protein misfolding tend to have primary fitness costs (see others in **figure S1**). **(D)** Summary of the Pearson correlations between protein stability and fitness for all 16 deep mutational scans. Correlations are closer to zero when fitness effects are collateral (light purple) and become stronger as the functional benefit of the protein increases (darker purple). The functional benefit of URA3 (involved in uracil production) was increased by removing uracil from media (Wu et al. 2022). The functional benefit of TEM1 (ampicillin degradation) was increased by adding ampicillin to the media (Stiffler et al. 2015).

We were surprised by this result, so as a control, we asked whether a mutation’s effect on stability could predict primary fitness costs. We observed a much stronger correlation between protein stability and primary fitness costs (**Fig 2C**; **Fig S1**). This correlation becomes stronger in environments where the primary fitness effects become more severe (**Fig 2D**), consistent with the idea that primary fitness costs are in part driven by misfolding-associated disruption of protein function. These results remain true when using other protein stability predictors, in addition to MutateX (**Fig S2–S3**) (Parthiban et al. 2006; Dehouck et al. 2009; Chen et al. 2020). These results, in line with prior literature (Yang et al. 2020; Lei et al. 2023; Küng et al. 2025), and supported by the Western data in **figure 3**, suggest that, on average, Mutate X is a reliable indicator of a mutation’s effect on protein stability. We therefore conclude that destabilizing mutations often underlie primary but not collateral fitness costs.

**Figure 3:**
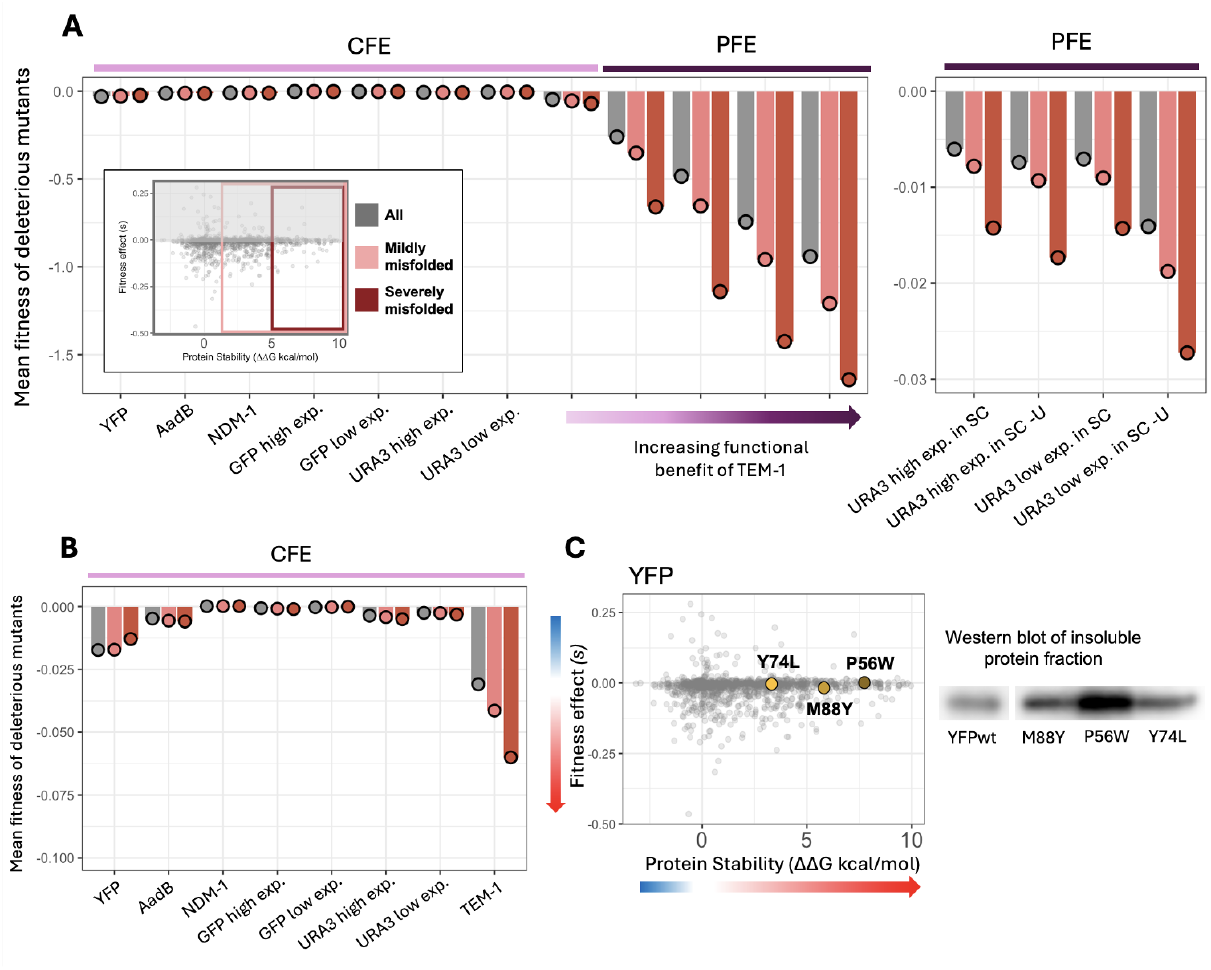
Mutations associated with severe misfolding do not have stronger collateral fitness costs than others. (**A**) In the inset, which is an example plot of collateral fitness effects versus stability, points on the far right tend to sit near zero on the vertical axis. This was unexpected because points on the far right represent protein variants that are predicted to be very destabilizing to protein folding. To understand the generality of this observation, we divided all protein variants into three groups: one representing the most severely destabilizing variants (red), another for variants with intermediate effects on stability (light red) and another comprising all variants (grey). Then we asked whether the average fitness cost of the variants in each group was different. For PFEs, we saw a clear trend whereby the fitness cost of the variants in each protein increases with the severity of misfolding. For CFEs, we saw no trends whereby the variants in one group had stronger fitness costs than others. Panel (**B**) displays this more clearly by showing the same data as in panel (**A**) with the vertical axis zoomed in. (**C**) Three of our misfolded YFP variants were previously studied in more detail (Quan et al. 2023) to confirm that they are more often found in the insoluble (misfolded) fraction of cells than wild type YFP. None of these variants have a measurable fitness cost, despite evidence from Western blots suggesting they are indeed misfolded..

### Misfolded proteins often have no collateral fitness cost in the conditions we survey

In all eight deep mutational scans for collateral fitness effects, we find some mutants that are predicted to be severely destabilizing, and yet are well tolerated. In fact, when we compare the average collateral fitness costs of the top destabilizing variants to all variants, we see no difference (**Fig 3A –3B**; red and grey points are at the same level on the vertical axis). On the other hand, the primary fitness costs of the top destabilizing variants are significantly larger (**Fig 3A**; red points are always lower than grey points on vertical axis). This comparison highlights that, even at the extremes, there is not a clear relationship between misfolding and collateral fitness costs.

We previously performed western blots on three misfolded YFP variants from our dataset: Y74L, M88Y, and P56W (Quan et al. 2023). These experiments demonstrated that each variant was enriched in the insoluble protein fraction (**Fig 3C**; Western Blots display insoluble fraction), confirming that they are misfolded. Both the western blot data and the MutateX prediction identify P56W as the most severely misfolded of the three, consistent with the structural disruption expected from replacing a small proline with a bulky tryptophan. Yet, the P56W variant, along with Y74L and M88Y, shows no measurable fitness cost compared to strains expressing wildtype YFP (**Fig 3C**; 3 yellow points fall at zero on the vertical axis). These data demonstrate that even severely misfolded proteins, when highly expressed, do not necessarily impair fitness. While prior studies have noted isolated examples of nontoxic misfolded proteins (Plata et al. 2010; Kafri et al. 2016), our results, spanning thousands of mutations across multiple proteins and organisms, suggest a broader need to re-evaluate whether misfolded protein toxicity may be less ubiquitous than previously believed.

### Collateral fitness costs often arise from mutations that do not hinder protein folding

Although the most destabilizing variants we study do not tend to have collateral fitness costs (**Fig 3)**, some other variants do indeed have large negative consequences on fitness. For example, in the AadB protein, the costliest variant is D143F, which results in a reproducible ∼40% reduction in growth rate compared to strains expressing wildtype AadB (Mehlhoff and Ostermeier 2023). Both Mutate X (**Fig 4**) and Western blots from previous studies indicate that this variant is not misfolded (Mehlhoff and Ostermeier 2023). Here we show that, across all 8 deep mutational scans of collateral fitness effects, the largest costs tend to arise from mutations that do not significantly hinder protein folding (**Fig 4A left**; variants with the lowest collateral fitness effects are found on the left side of the plots). On the other hand, the variants that result in the largest primary fitness costs are much more likely to be destabilizing (**Fig 4A right**; variants with the lowest primary fitness effects are found on the right side of the plots).

**Figure 4.**
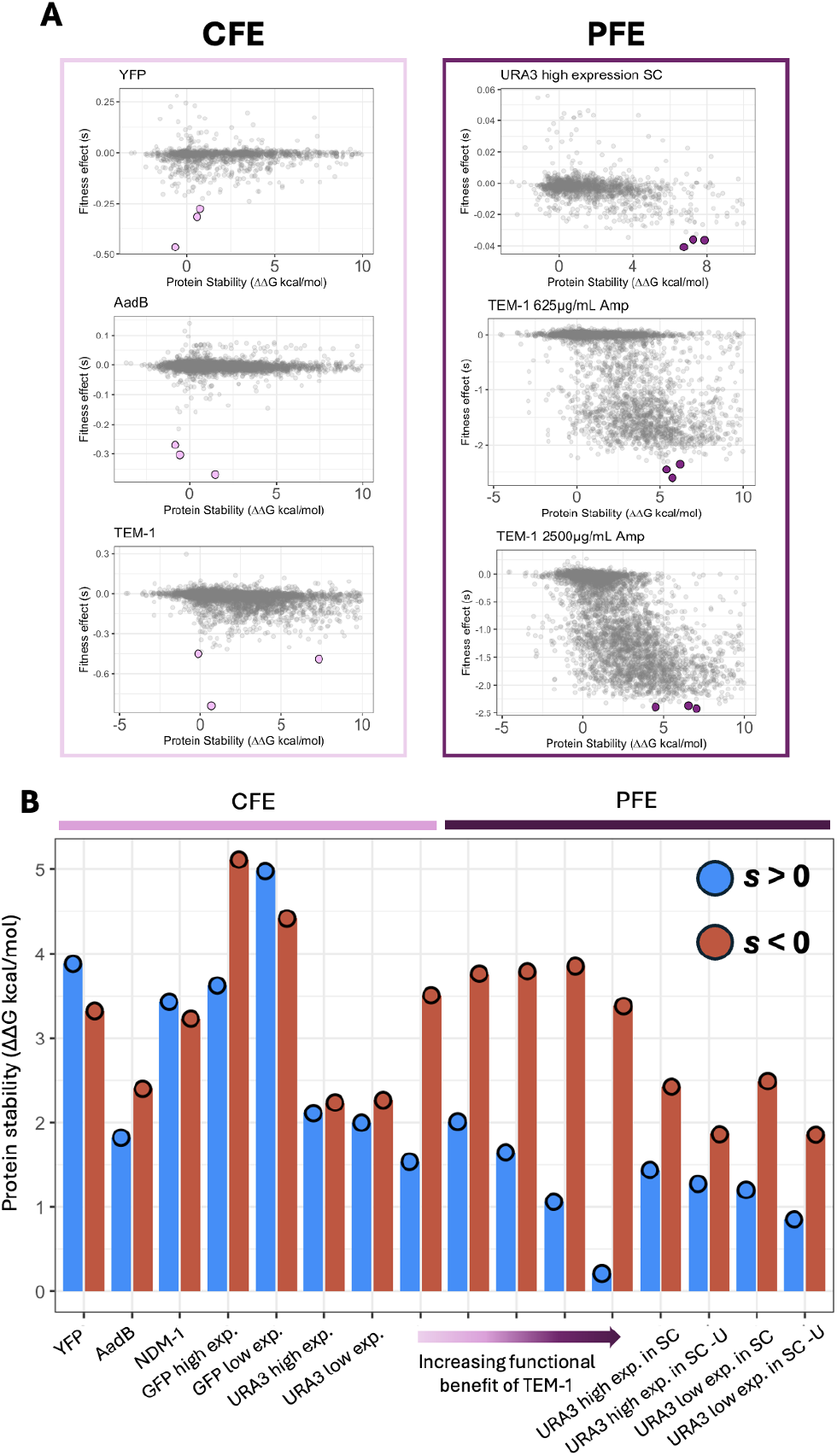
Deleterious mutants with collateral fitness effects are not often misfolded. (A) On the left: Example graphs showing that the most deleterious collateral fitness effects (dots highlighted in pink) generally are not severely misfolded. **On the right:** Example graphs showing that the mutants with most deleterious primary fitness effects (dots highlighted in purple) generally are severely misfolded. **(B)** Points showing the average ΔΔG value for mutants that are either deleterious *(s* < 0, red) or beneficial *(s* > 0, blue) for all proteins. For PFEs, the average ΔΔG for deleterious mutants is always greater than the ΔΔG for beneficial mutants, indicating that deleterious mutants are generally more unstable. For CFEs, there is no clear trend.

Moving beyond the costlier few variants for each protein, when we divide all variants into those that have negative fitness effects (s < 0) versus those that have positive fitness effects (s > 0), we see similar trends (or lack of trends for CFEs) (**Fig 4B**). Mutants that result in collateral fitness costs are no more likely to be destabilizing than are mutants that increase fitness (**Fig 4B**; left, red and blue bars are the same height). However, mutants that result in primary fitness costs tend to be far more destabilizing than those that do not (**Fig 4B**; right, red bars are taller than blue).

This finding begs questions about what causes collateral fitness costs. The cause of primary fitness costs is clear: loss of function due to protein misfolding. However, the cause of collateral fitness costs (typically arising from molecules that are not misfolded) is unclear. Previous work has confirmed both that: (1) a subset of the costliest mutations to AadB do not decrease protein stability, and (2) that they have strong effects on the transcriptome that are inconsistent with a transcriptional response to misfolding (Mehlhoff and Ostermeier 2023). This reinforces a major conclusion of our work, that collateral fitness costs are not routinely arising from protein misfolding.

## Materials and Methods

### Strains, growth conditions, and yeast transformation

All strains used in all experiments are listed in **[Supplementary Material S1]**.

Yeast culture and transformation were performed by the previously described methods (Amberg et al. 2005). A synthetic complete (SC) medium without uracil (-U) and/or histidine (-H) was used for yeast culture to maintain plasmids that utilize URA3 and HIS3 markers. Anhydrotetracycline (aTc) (Cayman Chemicals, 10009542) was prepared as a 0.2 mM stock solution in DMSO, diluted in DMSO, and added to the medium with the indicated aTc concentrations.

### Plasmids

All plasmids used in all experiments are listed in **[Supplementary Material S2]**.

### Barcoded YFP mutant library design and preparation

A library of guide/donor pairs was designed and ordered for CRISPR from Twist Bioscience (San Francisco, CA) following previous work (Sharon et al. 2018). This library included guide/donor pairs targeting 105 amino acid positions in YFP that were predicted by Dynamine, a protein structure prediction software, to have a high degree of rigidity (full guide/donor sequence list can be found in **Supplementary Material S3)**. These guide/donor pairs are also later used as barcodes for strain identification, and since many of these guide/donor pairs only differ by a single nucleotide, we shifted their length and position so that we could more easily distinguish them from each other. These guide/donor sequences were ligated into a modified version of the pZS165 yeast shuttle vector used in previous work (Sharon et al. 2018), generously shared by Shi-An Anderson and Hunter Fraser. This plasmid library was then transformed into a yeast strain possessing a Cas9. This yeast strain, called **YFP-CRISPR-WTC846**, also contains a modified copy of the YFP gene on its genome, of which expression is controlled by the WTC846 promoter (Azizoglu et al. 2019). The genotype of this strain is *MATα his3Δ1:: P*_*GAL1-GAL10-*_*SpCas9 P*_*GAL1-GAL10*_*-Ec86-RT leu2Δ0 met15Δ0:: P*_*RNR2*_*_TetR-NLS-TUP1 P*_*tetO7*.*1_*_*TetR-NLS ura3Δ0 ybr209wΔ::P*_*tetO7*.*1-*_*YFP:: HphMX6*. After transforming the guide/donor plasmid library into this yeast strain, gene editing via the CRISPEY method was performed by growing the strains in galactose (Sharon et al. 2018). PCR amplification of the YFP gene from a small subset of the engineered strains confirmed that all of the guide/donor sequences had successfully edited the YFP in this subset.

In the final YFP mutant library, the guide/donor pairs were sequenced at a depth of 10M reads to determine how many unique strains were present. In this experiment, out of all 1,995 strains, 1,502 strains were detected and successfully sequenced.

Spike-in control strains were also designed and constructed to be added to the final YFP mutant library directly before the competitive growth experiment. All plasmids are listed in **[Supplementary Material S2]**. 8 strains were constructed, named MU1-MU8. Mock-up sequences from MU1-MU4 were derived from the mCherry sequence, and mock-up sequences from MU5-MU8 were derived from the Cre recombinase sequence.

### Barcoded YFP mutant library competitive pooled growth assay

For the competitive growth assay, SC-HU + 2% Glucose media was prepared containing the various concentrations of aTc as described in **[Supplementary Material S4]**. Samples in each condition were prepared as described in **[Supplementary Material S4]**, which yielded a pool composed of 1% YFP wildtype (to use as a baseline) and the rest comprising YFP mutants. Pooled competitive growth was carried out in L-shaped glass tubes in an Advantec Bio-Photorecorder rocking incubator (Model TVS062CA). Each sample was cell-counted using a Beckman-Coulter Z-Series Cell Counter and diluted back every 24 h for a total of 8 rounds of growth, thus yielding 9 sample timepoints. Cells from each timepoint were frozen in 10% DMSO in both a cryotube for long term storage and a 1.5 mL tube for barcode extraction and sequencing.

### Plasmid extraction, PCR amplification of the barcode, and NGS sequencing

To calculate the relative frequencies of each strain and how they changed over time, the unique guide/donor region from each strain (its barcode) needed to be prepared for sequencing. To do so, each frozen yeast sample from each timepoint was thawed at RT and was centrifuged at 15,000 rpm for 1 min. After removal of the supernatant, 250 µL of yeast lysis solution 1 (0.1 M Na_2_EDTA, 1 M sorbitol, pH 7.5) and 1 µL of Zymolyase at 5U/µL (Zymo Research, E1005) were added to the pellet. The sample was incubated at 37°C for 30 min. After incubation, 250 µL of solution 2 (0.2 M NaOH, and 1% SDS) was added to the lysed sample and vortexed. Then, 250 µL of solution 3 (8.7% acetic acid and 5 M potassium acetate) was added and vortexed. After vortexing, the sample was centrifuged at 15,000 rpm for 10 min 750 µL of the supernatant was transferred to the spin column included in Monarch Plasmid Miniprep Kit (New England BioLabs, T1010L), and the column was centrifuged at 13,000 rpm for 1 min. After discarding the flow-through, 200 µL of Plasmid Wash Buffer 1 included in the kit was added to the column, and the column was centrifuged at 13,000 rpm for 1 min. After discarding the flow-through, 400 μL of Plasmid Wash Buffer 2 included in the kit was added to the column, and the column was centrifuged at 13,000 rpm for 1 min. After discarding the flow-through, the column was spun at 13,000 rpm for 1 min for the removal of wash buffer completely. The column was inserted into a new 1.5 mL tube, and 30 µL of DNA Elution Buffer included in the kit was added to the center of the matrix on the column. After waiting 1 min at room temperature, the tube was centrifuged at 13,000 rpm for 1 min to elute plasmids. The concentration of the plasmid was quantified by using Qubit dsDNA HS Assay Kit (Thermo Fisher Scientific, Q32854) on Qubit 4 Fluorometer (Thermo Fisher Scientific, Q33226).

PCR amplification of the barcode was performed by a two-step PCR scheme that controls for PCR duplicates as well as index swapping, very similar to the protocol described in (Levy et al. 2015; Kinsler et al. 2023). The indexes were selected by using BARCOSEL (Somervuo et al. 2018), a tool for selecting an optimal barcode, from the set of 288 barcodes prepared in (Levy et al. 2015) that allows for one nucleotide mismatch among the indexes. Another random 6-mer sequence was included in these primers to be used as a unique molecular identifier (UMI) to exclude PCR duplicates in downstream analysis. All primers in the first PCR were purified by HPLC to ensure the correct length. For the second PCR, IDT for Illumina DNA/RNA UD indexes SetA (Illumina, 20026121) and SetB (Illumina, 20026930) were used. All primers used in PCR amplification of the barcode were listed in **[Supplementary Material S5]**.

For the first step of the two-step PCR, the one reaction consisted of 13 µL of Nuclease free H2O, 10 µL of an extracted plasmid containing about 20 ng, 1 µL each of 10 µM forward and 10 µM reverse primer, and 25 µL of Hot Start Taq 2x Master Mix (New England BioLabs, M0496L). The first PCR was performed in hot-start PCR following the cycles: 1 cycle for 10 min at 94°C, 3 cycles for 3 min at 94°C; 1 min at 55°C; 1 min at 68°C, 1 cycle for 1 min at 68°C, and hold at 4°C. After the first PCR, the PCR product was cleaned up by using Monarch PCR and DNA Cleanup Kit (New England BioLabs, T1030L) following the manufacturer’s protocol, and the cleaned-up PCR product was eluted in 22 µL.

For the second step of the two-step PCR, the one reaction consisted of 14.5 µL of Nuclease free H2O, 20 µL of a cleaned-up PCR product, 10 µL of 5x Q5 Reaction Buffer, 2 µL each of forward and reverse primer of Illumina index primers, 1 µL of 10 mM dNTPs (Thermo Fisher Scientific, 18427088), 0.5 µL of Q5 Hot Start High-Fidelity DNA Polymerase (New England BioLabs, M0493L). The second PCR was performed in hot-start PCR following the cycles: 1 cycle for 30 s at 98°C, 2 cycles for 10 s at 98°C; for 20 s at 69°C; for 30 s at 72°C, 2 cycles for 10 s at 98°C; for 20 s at 67°C; for 30 s at 72°C, 20 cycles for 10 s at 98°C; for 20 s at 65°C; for 30 s at 72°C, 1 cycle for 3 min at 72°C, and hold at 4°C. The whole PCR product was loaded onto 2% of NuSieve 3:1 Agarose (LONZA, 50,090), and the band between 300 bp and 400 bp were sliced. The selected PCR product was extracted by Monarch DNA Gel Extraction Kit (New England BioLabs, T1020L) following the manufacturer’s protocol, and the extracted PCR product was eluted in 10 µL. The concentration of the product was quantified by using Qubit dsDNA HS Assay Kit on Qubit 4 Fluorometer.

The resulting samples were merged such that no two had similar Illumina or internal 8-mer indices, following a scheme to exclude any index swapping events that happened during NGS sequencing (Kinsler et al. 2020, 2023). Samples were sequenced on either a Novoseq or a Hiseq X to a coverage of an average 3.3 × 107 per sample. Since these amplicons libraries have low diversity, we spiked in 20% genomic DNA to all sequencing runs.

### Processing of NGS sequencing data

NGS sequencing data were demultiplexed into mate-pair files, a forward mate read1 (R1) file and a reverse mate read2 (R2) file, by Illumina sequencer software following an i5 and i7 indexes in an Illumina adaptor sequence. To exclude PCR duplicates in downstream processing, the UMIs of R1 and R2 files were extracted by using UMI-tools (Smith et al. 2017). with the following UMI-tools commands; umi_tools extract -I “R1 file” --bc-pattern = NNNNNN -S “extracted R1 output file” --read2-in = “R2 file” --bc-pattern2 = NNNNNN --read2-out = “extracted R2 output file.”

Then, the extracted R1 and R2 files were demultiplexed. We trimmed the 5′end region containing the index by using FLEXBAR (Roehr et al. 2017)(Dodt et al. 2012). with the following FLEXBAR commands; flexbar -r “extracted R1 file” -p “extracted R2 file” -b “index FASTA file for R1” -b2 “index FASTA file for R2” -bt LEFT -be 0.125 -n 10.

The STAR index files were generated from YFP reference sequences using the STAR aligner (Dobin et al. 2013) with the following STAR commands; STAR --runMode genomeGenerate --runThreadN 10 --genomeDir “STAR index output directory” --genomeFastaFiles “reference sequence FASTA file” --genomeSAindexNbases 8.

The reads in the demultiplexed R1 and R2 files were aligned to the STAR index sequences with the following STAR commands; STAR --genomeDir “STAR index output directory” --readFilesIn “demultiplexed R1 file” “demultiplexed R2 file” --runThreadN 10 --outSAMtype BAM Unsorted

--peOverlapNbasesMin 62 --peOverlapMMp 0 --outFilterMultimapNmax 1

--outFilterMismatchNmax 0 --alignEndsType EndToEnd --alignIntronMax 1 --alignIntronMin 2

--scoreDelOpen −10000 --scoreInsOpen −10000 --outFilterMatchNmin 137

--alignSoftClipAtReferenceEnds No --outReadsUnmapped Fastx.

The generated aligned sequence BAM file was sorted and indexed by using SAMtools (Li et al. 2009) with the following SAMtools commands; samtools sort -@ 8 -o “sorted output BAM file” “unsorted output BAM file,” samtools index “sorted BAM file.”

The duplicated reads in the indexed BAM file were excluded by using UMI-tools with the following UMI-tools commands; umi_tools dedup -I “indexed BAM file” --paired -S “output BAM file without duplicated reads” --chimeric-pairs = discard --unpaired-reads = discard --method cluster. Some small number of samples that received very high sequencing coverage took a very long time (days) to run using this method, presumably due to saturation of UMIs. Therefore we ran these using the same method but with the percentage rather than cluster method selected in the UMI-tools software.

The mapped reads in the BAM file without duplicated reads were counted by using SAMtools with the following SAMtools commands; samtools index “BAM file without duplicated reads,” samtools idxstats “indexed BAM file without duplicated reads” > “indexed SAM file without duplicated reads.”

### Calculation and analysis of protein misfolding using predictive algorithms

3D structures were taken from the Protein Data Bank (PDB accession codes are listed in [**Supplementary Material S6]**) and were run through the MutateX program (Tiberti et al. 2022) which is an automated version of FoldX (Guerois et al. 2002). The default additional running input files provided by MutateX upon download were used. The following MutateX command was used on all PDB files: mutatex “PDB file” --foldx-version=suite5 --foldx-binary= “FoldX binary file provided by MutateX” --nruns=5 --rotabase= “rotabase text file” &> “MutateX log output file”. Each MutateX output file that contained the calculated ΔΔG for each mutation for each protein was reformatted to include the selection coefficients associated with each mutation as well **[Supplementary Material S7]**.

The same PDB files for each protein were used for ΔΔG calculation and analysis using PoPMuSiC 2.1(Protein Mutant Stability Calculator) (Dehouck et al. 2011), CUPSAT (Cologne University Protein Stability Analysis Tool) (Parthiban et al. 2006), and PremPS (Chen et al. 2020). Each calculation for each protein was performed on the web server associated with each predictive algorithm.

### Data availability statement

The original contributions presented in this study are included in the article/Supplementary Materials, further inquiries can be directed to the corresponding author.

## Supplementary Figures

**Figure S1.**
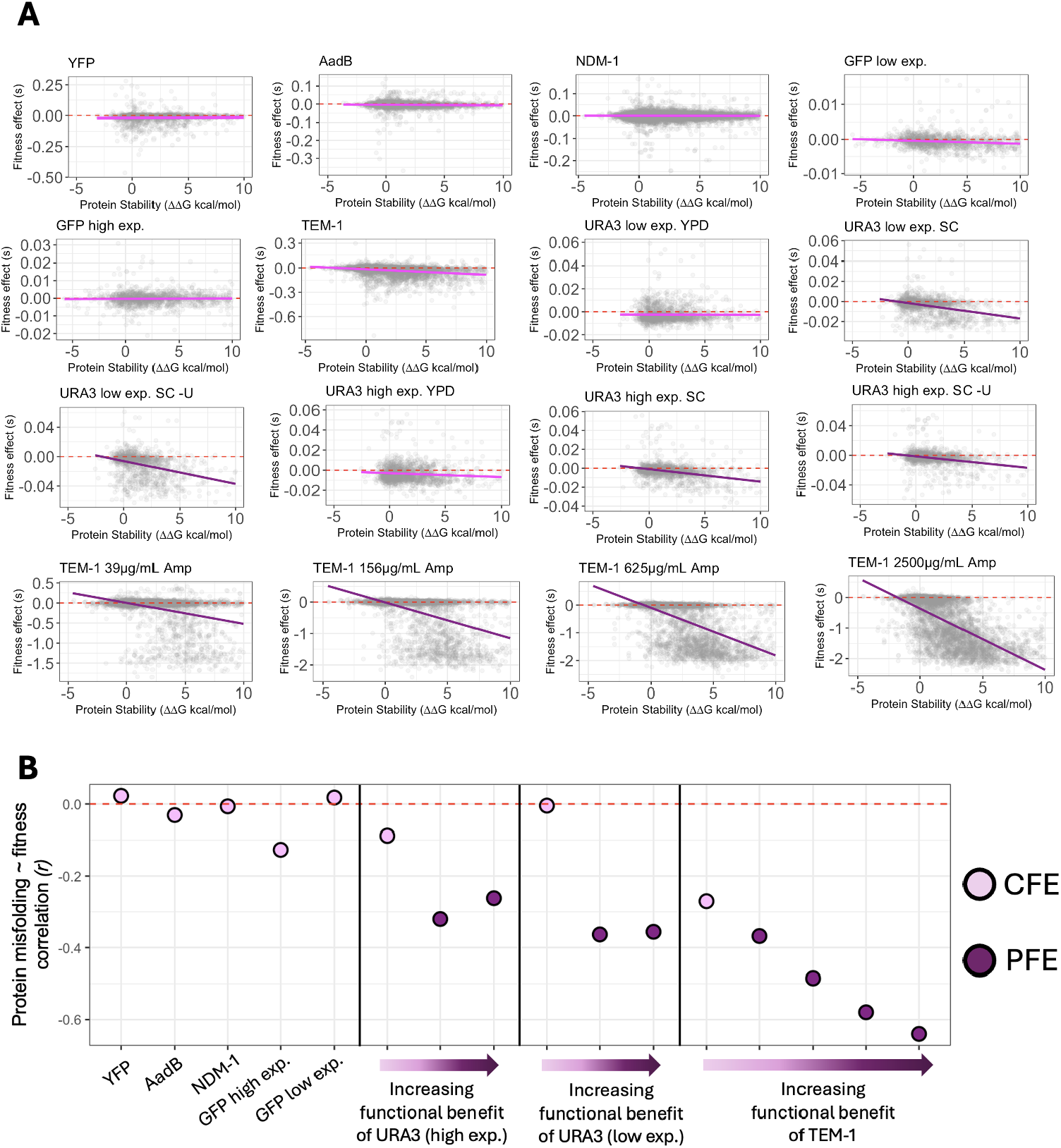
MutateX (FoldX) calculated ΔΔG values for each protein plotted against fitness *s*. This figure is similar to **Figure 3** in the main text but for all proteins.

**Figure S2:**
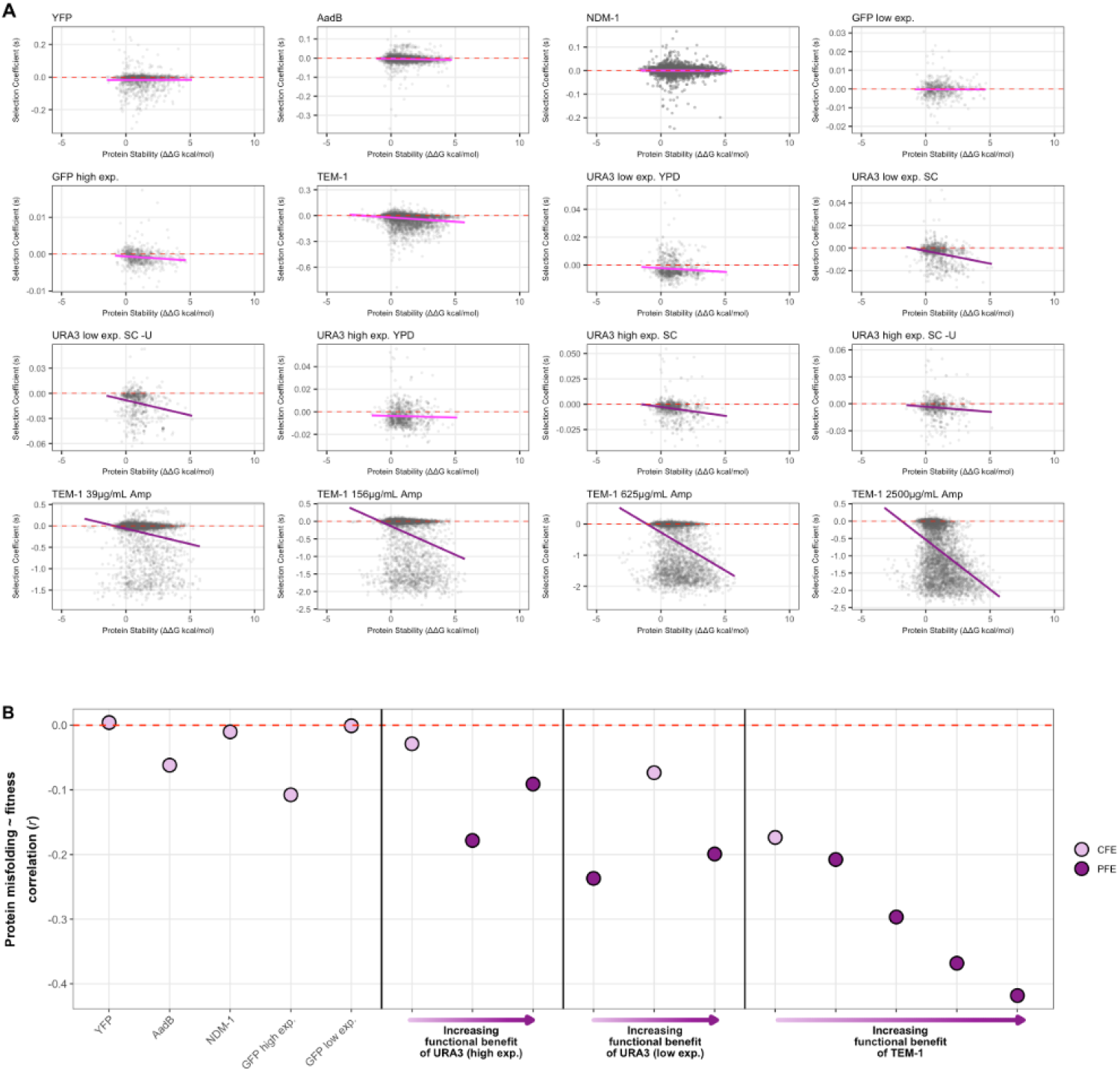
This figure shows the relationship between predicted protein stability and fitness effects when protein stability is calculated via PopMuSiC (Dehouck et al. 2011). The vertical axis is the selection coefficient, and the horizontal axis is the predicted change in protein stability. Every point represents a mutant. The line highlights the linear correlation between stability change and fitness. PopMuSiC estimate stability change by combining statistical potentials, amino acid volume terms, and a correction factor.

**Figure S3:**
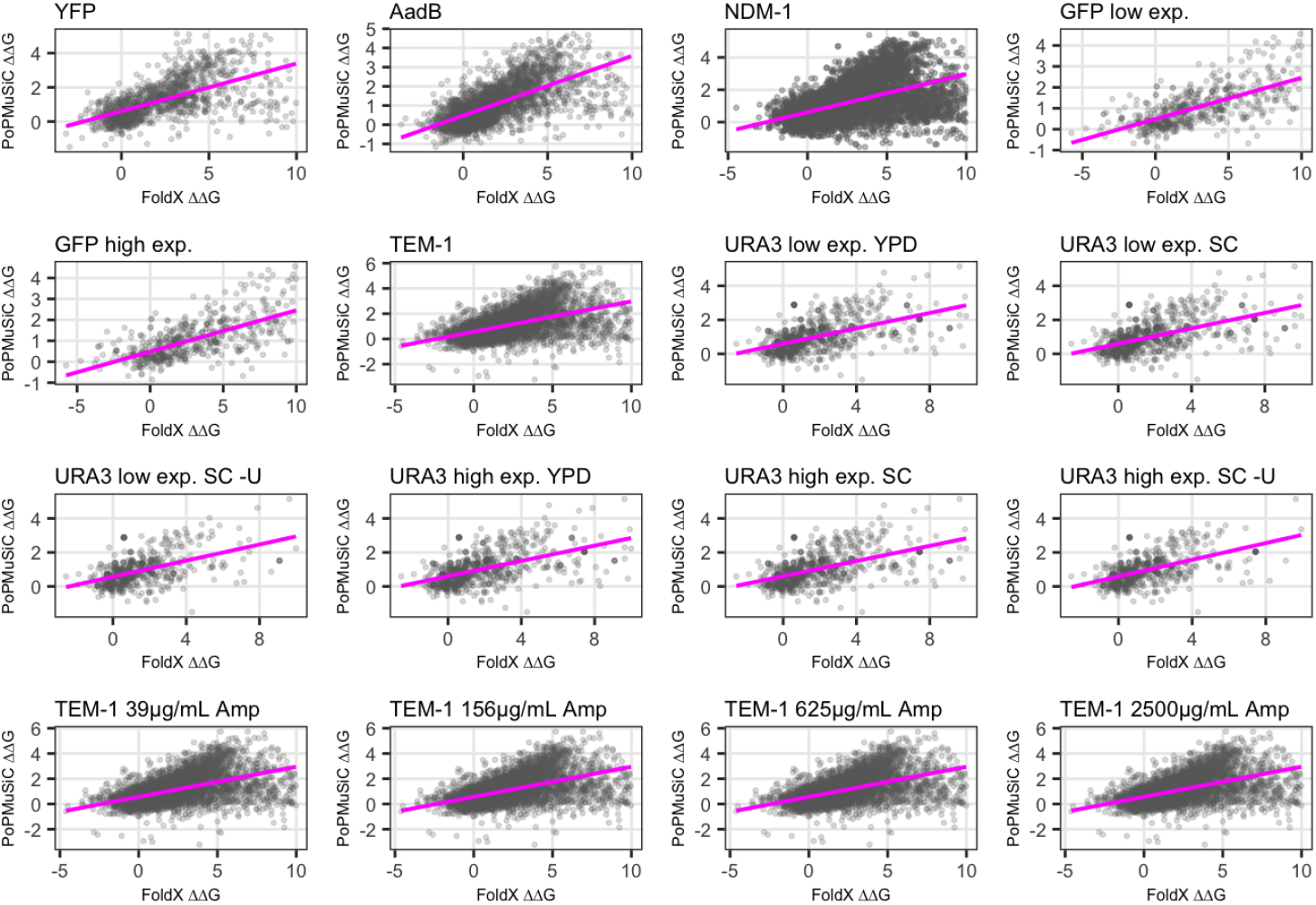
Comparison of FoldX and PoPMuSiC predictions. FoldX and PoPMuSiC are strongly correlated and generally yield similar stability estimates. Each panel shows one of the 16 datasets analyzed. Every point represents a single-site amino acid substitution, with the FoldX-predicted change in protein stability ΔΔG on the x-axis and the PoPMuSiC-predicted ΔΔG on the y-axis. The magenta line indicates the best-fit linear regression, highlighting the correlation between the two approaches.

## References

Abramov AY, Potapova EV, Dremin VV, Dunaev AV (2020) Interaction of Oxidative Stress and Misfolded Proteins in the Mechanism of Neurodegeneration. Life (Basel) 10.: 10.3390/life10070101

Alqahtani T, Deore S, Kide A, Ghosh A (2023) Mitochondrial dysfunction and oxidative stress in Alzheimer’s disease, and Parkinson’s disease, Huntington’s disease and Amyotrophic Lateral Sclerosis -An updated review. Mitochondrion 71:83–92

Amberg DC, Burke D, Strathern JN (2005) Methods in Yeast Genetics: A Cold Spring Harbor Laboratory Course Manual. CSHL Press

Azizoglu A, Brent R, Fabian R (2019) A precisely-titratable, variation-suppressed transcriptional controller to enable genetic discovery. bioRxiv

Azizoglu A, Brent R, Rudolf F (2021) A precisely adjustable, variation-suppressed eukaryotic transcriptional controller to enable genetic discovery. Elife 10.: 10.7554/eLife.69549

Bongioanni P, Del Carratore R, Corbianco S, et al (2021) Climate change and neurodegenerative diseases. Environ Res 201:111511

Brezic N, Gligorevic S, Sic A, Knezevic NN (2025) Protein Misfolding and Aggregation as a Mechanistic Link Between Chronic Pain and Neurodegenerative Diseases. Current Issues in Molecular Biology 47:259

Chen Y, Lu H, Zhang N, et al (2020) PremPS: Predicting the impact of missense mutations on protein stability. PLoS Comput Biol 16:e1008543

Dehouck Y, Grosfils A, Folch B, et al (2009) Fast and accurate predictions of protein stability changes upon mutations using statistical potentials and neural networks: PoPMuSiC-2.0. Bioinformatics 25:2537–2543

Dehouck Y, Kwasigroch JM, Gilis D, Rooman M (2011) PoPMuSiC 2.1: a web server for the estimation of protein stability changes upon mutation and sequence optimality. BMC Bioinformatics 12:151

Dobin A, Davis CA, Schlesinger F, et al (2013) STAR: ultrafast universal RNA-seq aligner. Bioinformatics 29:15–21

Dodt M, Roehr JT, Ahmed R, Dieterich C (2012) FLEXBAR-Flexible Barcode and Adapter Processing for Next-Generation Sequencing Platforms. Biology (Basel) 1:895–905

Drummond DA, Bloom JD, Adami C, et al (2005) Why highly expressed proteins evolve slowly. Proc Natl Acad Sci U S A 102:14338–14343

Drummond DA, Wilke CO (2008) Mistranslation-induced protein misfolding as a dominant constraint on coding-sequence evolution. Cell 134:341–352

Echave J, Wilke CO (2017) Biophysical Models of Protein Evolution: Understanding the Patterns of Evolutionary Sequence Divergence. Annu Rev Biophys 46:85–103

Eguchi Y, Bilolikar G, Geiler-Samerotte K (2019) Why and how to study genetic changes with context-dependent effects. Curr Opin Genet Dev 58-59:95–102

Eguchi Y, Makanae K, Hasunuma T, et al (2018) Estimating the protein burden limit of yeast cells by measuring the expression limits of glycolytic proteins. Elife 7.: 10.7554/eLife.34595

Geiler-Samerotte KA, Dion MF, Budnik BA, et al (2011) Misfolded proteins impose a dosage-dependent fitness cost and trigger a cytosolic unfolded protein response in yeast. Proc Natl Acad Sci U S A 108:680–685

Geiler-Samerotte KA, Hashimoto T, Dion MF, et al (2013) Quantifying condition-dependent intracellular protein levels enables high-precision fitness estimates. PLoS One 8:e75320

Gidalevitz T, Krupinski T, Garcia S, Morimoto RI (2009) Destabilizing protein polymorphisms in the genetic background direct phenotypic expression of mutant SOD1 toxicity. PLoS Genet 5:e1000399

Gonzalez-Garcia M, Fusco G, De Simone A (2021) Membrane Interactions and Toxicity by Misfolded Protein Oligomers. Front Cell Dev Biol 9:642623

Guerois R, Nielsen JE, Serrano L (2002) Predicting changes in the stability of proteins and protein complexes: a study of more than 1000 mutations. J Mol Biol 320:369–387

Hartl FU (2017) Protein Misfolding Diseases. Annu Rev Biochem 86:21–26

Kafri M, Metzl-Raz E, Jona G, Barkai N (2016) The Cost of Protein Production. Cell Rep 14:22–31

Karapanagioti F, Atlason ÚÁ, Slotboom DJ, et al (2024) Fitness landscape of substrate-adaptive mutations in evolved amino acid-polyamine-organocation transporters. Elife 13.: 10.7554/eLife.93971

Kinsler G, Geiler-Samerotte K, Petrov DA (2020) Fitness variation across subtle environmental perturbations reveals local modularity and global pleiotropy of adaptation. Elife 9.: 10.7554/eLife.61271

Kinsler G, Schmidlin K, Newell D, et al (2023) Extreme sensitivity of fitness to environmental conditions: Lessons from #1BigBatch. J Mol Evol 91:293–310

Kintaka R, Makanae K, Moriya H (2016) Cellular growth defects triggered by an overload of protein localization processes. Sci Rep 6:31774

Küng C, Protsenko O, Vanella R, Nash MA (2025) Deep mutational scanning reveals a de novo disulfide bond and combinatorial mutations for engineering thermostable myoglobin. Protein Science 34:e70112

Lei R, Hernandez Garcia A, Tan TJC, et al (2023) Mutational fitness landscape of human influenza H3N2 neuraminidase. Cell Rep 42:113356

Levy SF, Blundell JR, Venkataram S, et al (2015) Quantitative evolutionary dynamics using high-resolution lineage tracking. Nature 519:181–186

Li H, Handsaker B, Wysoker A, et al (2009) The Sequence Alignment/Map format and SAMtools. Bioinformatics 25:2078–2079

McFarland CD, Korolev KS, Kryukov GV, et al (2013) Impact of deleterious passenger mutations on cancer progression. Proc Natl Acad Sci U S A 110:2910–2915

McFarland CD, Yaglom JA, Wojtkowiak JW, et al (2017) The Damaging Effect of Passenger Mutations on Cancer Progression. Cancer Res 77:4763–4772

Mehlhoff JD, Ostermeier M (2023) Genes Vary Greatly in Their Propensity for Collateral Fitness Effects of Mutations. Mol Biol Evol 40.: 10.1093/molbev/msad038

Mehlhoff JD, Stearns FW, Rohm D, et al (2020) Collateral fitness effects of mutations. Proc Natl Acad Sci U S A 117:11597–11607

Morimoto RI, Cuervo AM (2014) Proteostasis and the aging proteome in health and disease. J Gerontol A Biol Sci Med Sci 69 Suppl 1:S33–8

Oromendia AB, Dodgson SE, Amon A (2012) Aneuploidy causes proteotoxic stress in yeast. Genes Dev 26:2696–2708

Pakula AA, Sauer RT (1989) Genetic analysis of protein stability and function. Annu Rev Genet 23:289–310

Pál C, Papp B, Hurst LD (2001) Highly expressed genes in yeast evolve slowly. Genetics 158:927–931

Pancotti C, Benevenuta S, Birolo G, et al (2022) Predicting protein stability changes upon single-point mutation: a thorough comparison of the available tools on a new dataset. Brief Bioinform 23.: 10.1093/bib/bbab555

Parthiban V, Gromiha MM, Schomburg D (2006) CUPSAT: prediction of protein stability upon point mutations. Nucleic Acids Res 34:W239–42

Plata G, Gottesman ME, Vitkup D (2010) The rate of the molecular clock and the cost of gratuitous protein synthesis. Genome Biol 11:R98

Quan N, Eguchi Y, Geiler-Samerotte K (2023) Intra-FCY1: a novel system to identify mutations that cause protein misfolding. Front Genet 14:1198203

Roehr JT, Dieterich C, Reinert K (2017) Flexbar 3.0 - SIMD and multicore parallelization. Bioinformatics 33:2941–2942

Santra M, Dill KA, de Graff AMR (2019) Proteostasis collapse is a driver of cell aging and death. Proc Natl Acad Sci U S A 116:22173–22178

Serohijos AWR, Rimas Z, Shakhnovich EI (2012) Protein biophysics explains why highly abundant proteins evolve slowly. Cell Rep 2:249–256

Sharon E, Chen S-AA, Khosla NM, et al (2018) Functional Genetic Variants Revealed by Massively Parallel Precise Genome Editing. Cell 175:544–557.e16

Sisodiya SM, Gulcebi MI, Fortunato F, et al (2024) Climate change and disorders of the nervous system. Lancet Neurol 23:636–648

Smith T, Heger A, Sudbery I (2017) UMI-tools: modeling sequencing errors in Unique Molecular Identifiers to improve quantification accuracy. Genome Res 27:491–499

Somervuo P, Koskinen P, Mei P, et al (2018) BARCOSEL: a tool for selecting an optimal barcode set for high-throughput sequencing. BMC Bioinformatics 19:257

Soto C, Pritzkow S (2018) Protein misfolding, aggregation, and conformational strains in neurodegenerative diseases. Nat Neurosci 21:1332–1340

Stiffler MA, Hekstra DR, Ranganathan R (2015) Evolvability as a function of purifying selection in TEM-1 β-lactamase. Cell 160:882–892

Tiberti M, Terkelsen T, Degn K, et al (2022) MutateX: an automated pipeline for in silico saturation mutagenesis of protein structures and structural ensembles. Brief Bioinform 23.: 10.1093/bib/bbac074

Tilk S, Frydman J, Curtis C, Petrov DA (2024) Cancers adapt to their mutational load by buffering protein misfolding stress. Elife 12.: 10.7554/eLife.87301

Tokuriki N, Stricher F, Schymkowitz J, et al (2007) The stability effects of protein mutations appear to be universally distributed. J Mol Biol 369:1318–1332

Wu SA, Li ZJ, Qi L (2025) Endoplasmic reticulum (ER) protein degradation by ER-associated degradation and ER-phagy. Trends Cell Biol 35:576–591

Wu Z, Cai X, Zhang X, et al (2022) Expression level is a major modifier of the fitness landscape of a protein coding gene. Nat Ecol Evol 6:103–115

Yang J, Naik N, Patel JS, et al (2020) Predicting the viability of beta-lactamase: How folding and binding free energies correlate with beta-lactamase fitness. PLoS One 15:e0233509

Yang J-R, Liao B-Y, Zhuang S-M, Zhang J (2012) Protein misinteraction avoidance causes highly expressed proteins to evolve slowly. Proc Natl Acad Sci U S A 109:E831–40

Zhang J, Yang J-R (2015) Determinants of the rate of protein sequence evolution. Nat Rev Genet 16:409–420

